# Automation Disrupts, Explanations Restore: The Neural Signatures of Agency Loss and Recovery in Human–AI Interaction

**DOI:** 10.64898/2026.07.22.740020

**Authors:** Eléonore Houdoyer, Solène Le Bars, Valérian Chambon

## Abstract

Automation has been shown to weaken the sense of agency (SoA), the experience of controlling one’s actions and their outcomes, by disrupting the predictive link between intention and effect. Explainable AI (XAI) has been proposed as a solution, yet the neurocognitive mechanisms through which explanations restore agency remain unclear. Across three EEG experiments using an autonomous-driving paradigm, we examined how automation and different forms of AI explanations modulate explicit agency judgments and early neural markers of agency-related predictive processing. In Experiment 1, automation reduced explicit feelings of control and was associated with reduced sensory attenuation, as reflected by increased P1–N1 amplitudes, decreased N1–P2 amplitudes, and delayed N1 latencies. In Experiment 2, distal (goal-level) explanations partially restored agency and selectively modulated early auditory responses, decreasing P1–N1 and increasing N1–P2 amplitudes. In Experiment 3, combining distal and proximal (trajectory-level) explanations produced the strongest behavioural and neural restoration of agency, yielding a graded attenuation of P1–N1 and enhanced N1–P2 responses along with accelerated N1 latencies. Across all experiments, mismatch negativity (MMN) remained unaffected, indicating that pre-attentive deviance detection is preserved regardless of agency or explainability. Together, these results identify component-specific EEG markers that track fluctuations in the sense of agency and demonstrate that multi-level intention sharing by AI systems enhances both predictive engagement and explicit control experience. This work provides a neurocognitive foundation for designing explainable autonomous systems capable of maintaining user agency.

## 1. Introduction

The sense of agency (SoA) refers to the subjective experience of initiating and controlling one’s own actions and their outcomes. This self-attributed control is fundamental not only to motor coordination but also to the attribution of causal, legal, and moral responsibility (Chambon & Bigenwald, 2019). As intelligent systems increasingly mediate human decisions and actions, a central question arises: do individuals still experience agency, and therefore accountability, when outcomes are generated or executed by artificial agents?

Recent research in human-AI interaction (HAI) indicates that automation can profoundly modulate the sense of agency (SoA). When humans shift from direct control to supervisory roles, SoA typically declines, a phenomenon associated with the out-of-the-loop performance problem, marked by reduced vigilance, impaired error detection, and diminished engagement (Endsley & Kiris, 1995; Christoffersen & Woods, 2002; Sarter et al., 1997). However, this attenuation is not uniform. In some contexts, automation can even enhance perceived control, suggesting that agency in automated environments is context-dependent and shaped by specific system design features (Berberian et al., 2012; Vantrepotte et al., 2022).

A key determinant of SoA is the ability to form, anticipate, and represent intentions. Human intentions are hierarchically organized, ranging from motor intentions that specify fine-grained kinematic parameters, to proximal intentions that guide near-term subgoals, to distal intentions that encode abstract, long-term objectives (Pacherie, 2008; Chambon et al., 2011, 2017). This hierarchical architecture supports predictive processing of action outcomes and facilitates the dynamic integration of feedback during control. In joint human action, co-representing a partner’s layered intentions facilitates coordination, anticipation, and mutual prediction (Le Bars et al., 2022). By contrast, when interacting with artificial agents, this capacity for shared intentional representation is hindered by the opacity of algorithmic decision-making. Such opacity limits users’ ability to infer the agent’s goals, thereby weakening both their SoA and their trust (Pagliari et al., 2022; Liao et al., 2020).

To address these challenges, eXplainable AI (XAI) frameworks aim to make artificial systems more interpretable and predictable to human users (Putnam & Conati, 2019). Yet the specific forms of explanation that most effectively restore transparency and a sense of control remain debated (Vantrepotte et al., 2022). Evidence from joint action research suggests that intention sharing plays a critical role in supporting agency: when co-actors explicitly communicate their goals, partners experience stronger coordination, predictive alignment, and mutual understanding (van der Wel et al., 2012; Le Bars et al., 2020, 2022). Extending this principle to human–AI interaction, enabling artificial agents to disclose their proximal (path-level) and distal (goal-level) intentions may provide users with the predictive cues necessary to sustain agency, transparency, and trust.

In a previous behavioral study using an automated driving paradigm (Houdoyer et al., under review), we found that explicit (subjective) agency ratings increased across all explanation types, proximal, distal, and combined, relative to the no-explanation condition. By contrast, the implicit measure of agency (Intentional Binding, IB), referring to the perceived temporal compression between an action and its effect (see Moore et al., 2012), was selectively enhanced under combined explanations, whereas proximal or distal cues alone had no effect. These findings suggest that while any form of explanation can strengthen explicit judgments of control, only dual-level (proximal + distal) intention sharing effectively supports the implicit predictive mechanisms that contribute to the SoA. However, both explicit ratings and intentional binding measures present inherent limitations for studying agency in ecologically valid contexts. In real-world scenarios, such as autonomous driving, participants cannot perform time-estimation tasks without disrupting performance, and subjective ratings remain vulnerable to introspective and demand-related biases. These constraints underscore the need for objective, non-invasive approaches capable of capturing the SoA in real time and under ecologically valid conditions, without interfering with ongoing behavior.

Neurophysiological markers provide a promising way to overcome these limitations by offering direct access to the brain mechanisms underlying action monitoring and predictive control (see Haggard, 2017, for a review). Among available techniques, electroencephalography (EEG) is particularly well suited to this purpose because it combines millisecond-level temporal resolution with continuous, non-invasive recording of neural activity. It therefore allows the investigation of the fast sensory and predictive processes that contribute to the experience of control.

Within this framework, most EEG studies of agency and voluntary action have focused on event-related potentials (ERPs) elicited by the sensory consequences of action. These components index early sensory encoding and prediction-error processing, particularly within the auditory modality through components such as the N1 and P2, as well as mismatch-related responses including the mismatch negativity (MMN) and the N2b (Baess et al., 2011; Hughes et al., 2013; Kühn et al., 2011; Le Bars et al., 2019).

Under conditions associated with stronger agency, self-generated or well-predicted outcomes typically elicit attenuated N1 amplitudes (approximately 80–130 ms over fronto-central regions), reflecting predictive suppression when sensory consequences match internal action-related predictions (Baess et al., 2011; Hughes et al., 2013; Timm et al., 2014; Sugimoto, Kimura, & Takeda, 2021). The P2 component (approximately 150–250 ms) has also been linked to agency-related processing, although its modulation is less consistent across studies than that of the N1. Depending on task structure and interpretative demands, stronger agency has been associated with either reduced or enhanced P2 amplitudes (Timm et al., 2016; Pinheiro et al, 2019; Bolt and Loehr et al., 2021; Han et al.,2021; Hughes et al.,2025). This variability suggests that the P2 does not constitute a straightforward marker of sensory attenuation (see Egan et al., 2026), but rather reflects later evaluative or integrative stages of action–outcome processing (Han et al., 2021; Hughes et al., 2025).

Importantly, numerous studies in action-related auditory processing suggest that sensory attenuation does not imply the isolated suppression of a single ERP component, but instead reflects a reconfiguration of early successive components within the P1–N1–P2 complex (Timm et al., 2014; Ross et al., 2017; Korka et al., 2021). Accordingly, peak-to-peak measures were used in the present study to assess changes in sensory attenuation arising from action–outcome prediction. This approach allows the capture of condition-dependent modulations of early sensory processing while reducing ambiguity related to baseline fluctuations and temporal overlap between ERP components, especially in complex and naturalistic paradigms.

Building on these findings, the present study aimed to identify neurophysiological correlates associated with the SoA under varying levels of automation and AI explainability. Specifically, we examined whether behavioural effects previously observed in human–AI interaction, namely the reduction of explicit agency under automation (Berberian et al.,2012) and its partial restoration through AI explanations (Vantrepotte et al., 2022; Houdoyer et al., under review), would also be detectable at the neural level.

To address this question, explicit measure of agency (i.e., subjective ratings of perceived control) were combined with electrophysiological indices of predictive processing within a sensory attenuation paradigm. This approach allows us to assess whether the same experimental manipulations modulate neural markers associated with action–outcome prediction and sensory processing.

Notably, we hypothesized that voluntary motor control, relative to automation, would be associated with reduced amplitudes of early auditory ERP complexes, as reflected by attenuated P1–N1 peak-to-peak measures. Such modulation would be consistent with stronger predictive attenuation when action outcomes are generated by one’s own motor system. Conversely, automation was expected to weaken action–outcome predictive coupling, leading to increased early sensory responses. We further hypothesized that automation, relative to voluntary motor control, would alter the configuration of the auditory ERP complex spanning the N1–P2 interval. Because previous studies have reported both increases and decreases of P2-related activity depending on task demands and evaluative context, we did not formulate a directional hypothesis regarding N1–P2 modulation.

Considering the impact of AI explanations, we expected that explanatory information would modulate early auditory neural signatures by improving the predictability of interaction outcomes. Distal explanations, which provide information about the system’s high-level goals, were expected to partially restore predictive processing, as it should be objectivized via P1–N1 attenuation and N1–P2 modulation. Finally, combined proximal and distal explanations, integrating both action-level and goal-level information in line with hierarchical models of intention (Pacherie, 2008), were predicted to produce the strongest effects on early auditory components. By providing a more complete representation of the system’s behaviour, such explanations may enhance predictive coherence, leading to increased sensory attenuation and reduced prediction-error responses (Widmann et al., 2022). This pattern would result in neural responses more closely resembling those observed for self-generated actions, thereby indicating a more complete restoration of the SoA.

Complementing the analysis of early auditory ERP complexes, the MMN (approximately 100–250 ms, fronto-central) provides a pre-attentive index of auditory prediction-error processing: its amplitude increases when sensory feedback violates established expectations and decreases when outcomes are predictable (Näätänen et al., 2011). Although not a direct marker of agency per se, the MMN is informative about one of its core computational mechanisms, namely the comparison between expected and actual outcomes. Accordingly, MMN was analyzed to explore whether deviant auditory consequences were differentially processed when generated by voluntary actions versus human–AI interactions. Together, early auditory ERP complexes (P1–N1–P2) and MMN provide temporally precise neural markers of how predictive and feedback-related mechanisms are modulated by automation, explainability, and the degree of intentional control in human–AI interaction.

To test our hypotheses, we conducted three EEG experiments designed to systematically examine how automation and AI explainability modulate the neurophysiological correlates associated with the SoA. Across experiments, participants engaged in a simplified driving task in which they either actively controlled a car to reach a target or monitored an AI-controlled vehicle performing the same goal-directed action (Experiment 1). Depending on the experimental condition, the AI provided either no explanation or verbal information about its intended actions (Experiments 2 and 3). To ensure high-quality EEG recordings, the original behavioral paradigm (Houdoyer et al., under review) was simplified to reduce motor and cognitive demands.

In Experiments 1 and 2, each navigational strategy was represented by a single target, and the car executed only one brief movement per trial, an adaptation intended to minimize cognitive load and help ensure that neural responses more closely reflected agency-related processes. In Experiment 3, the task was extended to include two targets per strategy, enabling the examination of combined proximal and distal explanations. This controlled increase in task complexity reintroduced path-level information while maintaining experimental precision, allowing a direct comparison between distal-only and dual-level (proximal + distal) intention sharing. This design choice was guided by our previous behavioral findings, which showed that dual-level explanations produced the strongest enhancement of agency. Across all experiments, we analyzed early auditory-evoked complexes (P1–N1, N1–P2) and the mismatch negativity (MMN) as neural indices of sensory attenuation and predictive processing. In addition, explicit agency ratings were collected on 20% of trials as behavioral controls, enabling verification of whether neural modulations of agency aligned with participants’ subjective sense of control.

## 2. Methods

### 2.1 Participants

EEG data were collected from 87 paid volunteers, who were assigned to three independent groups of 29 participants for Experiments 1, 2, and 3. All participants reported normal hearing, normal or corrected-to-normal vision, no colour blindness, and no history of neurological or psychiatric disorders.

Thirty participants were excluded from the analyses: 20 because of excessive EEG noise, mainly prolonged low-frequency drifts that led to the rejection of approximately half of the epochs and left fewer than 40 usable mismatch trials, and 10 because of technical problems, including trigger-recording failures and EEG-cap malfunctions. The final sample therefore comprised 20 participants in Experiment 1 (10 females, 10 males; mean age = 23.57 ± 3.34 years), 18 participants in Experiment 2 (8 females, 10 males; mean age = 23.07 ± 3.26 years), and 19 participants in Experiment 3 (10 females, 9 males; mean age = 23.68 ± 3.21 years). All participants were naïve to the purpose of the study and provided written informed consent before taking part. They received a fixed compensation of €15 and were informed that they could receive an additional performance-based bonus if their score ranked among the three highest. The exact amount of the bonus was not specified beforehand. At the end of the experiment, all participants received an additional €5, resulting in a total compensation of €20, and were debriefed about the bonus procedure. The study was approved by the local ethics committee (IRB00003888) and conducted in accordance with the Declaration of Helsinki (1964, revised 2013).

### 2.2 Materials

All three experiments were conducted in a sound-attenuated booth at the École Normale Supérieure. The task was implemented in PsychoPy (version 2025.1.1) with PyLSL (version 1.14) for event triggering, and presented in full-screen mode on a 14-inch LCD monitor (1920 × 1080 pixels, 60 Hz), positioned approximately 60 cm from the participant against a uniform grey background (RGB: 128, 128, 128). Auditory feedback was delivered through Sennheiser HD 280 Pro closed-back headphones, calibrated to 80 dB SPL. Participant responses were recorded via a standard computer keyboard.

### 2.3 Experimental Design, Stimuli, and Procedure

Each trial began with a grey background displaying a car icon and target icons corresponding to two distinct navigational strategies designed to manipulate distal-level intentionality, defined as long-term goal selection. A red heart indicated the safe strategy, whereas a banknote symbol indicated the risky strategy. The safe strategy was characterised by lower risk and lower reward, whereas the risky strategy was characterised by higher risk and higher reward. Specifically, the safe strategy entailed a 20% obstacle probability and yielded a 2.5-point reward, whereas the risky strategy involved a 50% obstacle probability and offered a 4-point reward.

In the Motor/no-AI condition, these strategy characteristics were fully implemented, allowing participants to familiarise themselves with the two strategies and their associated features, namely speed, risk, and reward. This phase was also intended to establish prior beliefs about the consequences associated with each strategy. In the belief-based AI conditions, by contrast, strategy-related differences in obstacle probability were removed to ensure experimental control. Obstacle probability was fixed at 10% for both strategies, while reward magnitude continued to vary according to the selected strategy. This low and constant obstacle rate was intended to promote trust in the purported AI-based assistance system and to minimise data loss. Obstacle trials were included to maintain participants’ attention and preserve the plausibility of the reward–risk structure of the driving task. However, they were not treated as trials of interest and were excluded from the main analyses.

The spatial layout of the targets differed across experiments. In Experiments 1 and 2, two targets, one per strategy, were presented equidistantly to the left and right of the car to maintain a simple, symmetrical visual layout (see Figures 1A and 1B). In Experiment 3, four targets, two per strategy, were positioned in the four corners of the screen, introducing two possible trajectories per strategy (see Figures 1C and 1D). This configuration allowed proximal-level intentionality, defined as path-level action selection, to be manipulated in addition to distal-level intentionality, defined as strategy selection.

**Figure 1.**
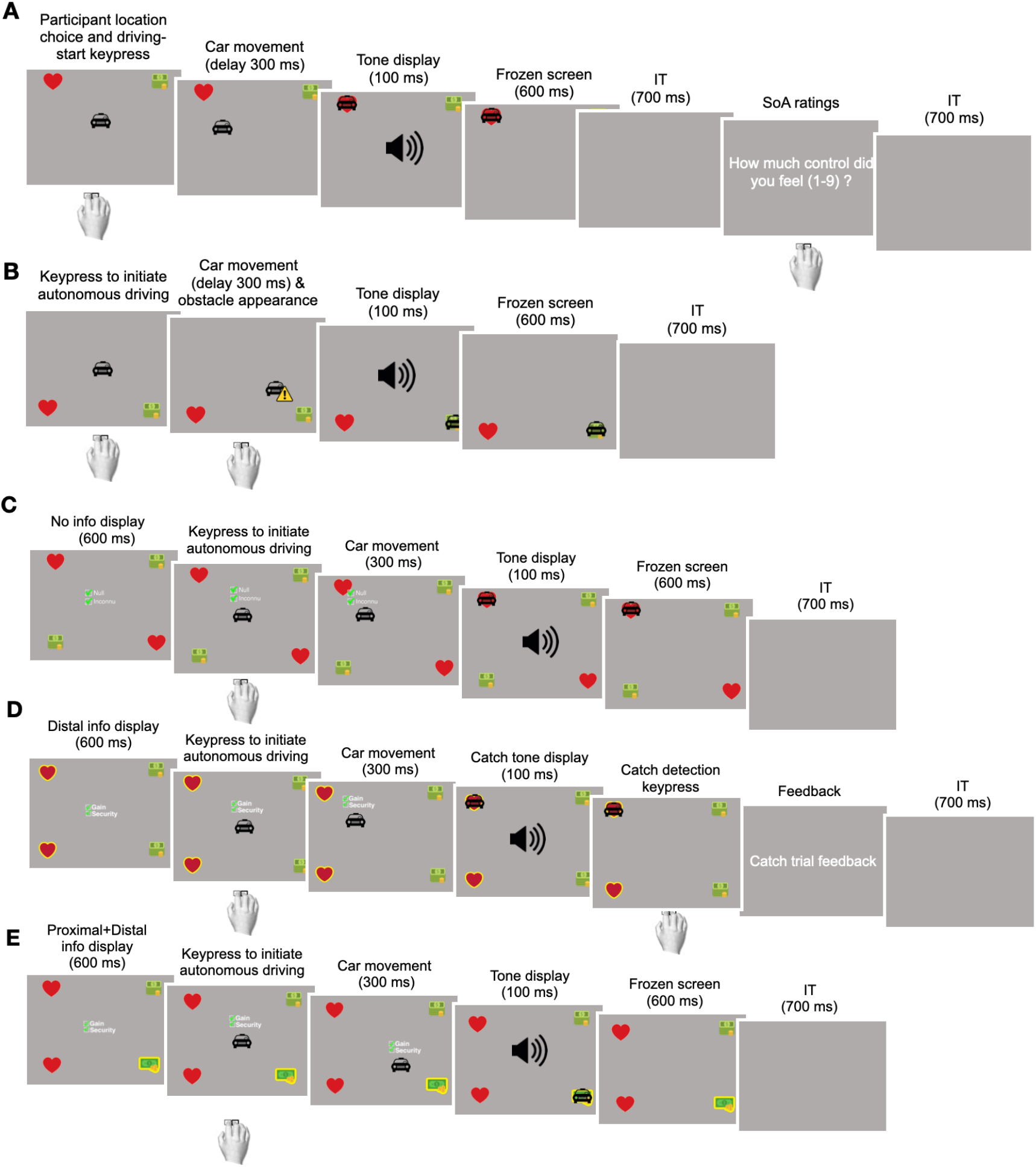
Task layout and trial timeline across automation and AI explanation conditions. Each trial began with the presentation of target(s) defining the available strategy options (Experiments 1-2: two targets; Experiment 3: four targets, two per strategy) and their associated trajectories. In the motor condition (A), participants selected the target using the left or right arrow key. In AI conditions (B-E), participants initiated the trial by pressing the space bar, while the AI selected the strategy and trajectory. The car then moved toward the chosen target for 300 ms. In a subset of trials, an obstacle (yellow warning panel) appeared during movement, requiring a rapid response. Upon target arrival, an auditory tone was presented (match ≈80%, mismatch/oddball ≈20%), and in 20% of trials, participants rated their sense of agency on a 9-point scale. In Experiments 2 and 3, AI conditions included visual explanation cues displayed for 600 ms before movement onset. The no-explanation condition (C) presented green checkmarks with meaningless placeholder text. In the distal explanation condition (D), the AI-selected strategy was indicated by highlighting the corresponding targets and accompanied by green checkmarks and explanatory text. In the combined proximal-and-distal explanation condition (E), the specific trajectory was additionally indicated by highlighting a single target among the two available for the selected strategy.

After target presentation, either the participant or the purported AI system selected the strategy and, when applicable, the trajectory. In the Motor/no-AI condition, participants selected and initiated their chosen strategy by pressing either the left or right arrow key, depending on the side of the screen on which their preferred target appeared. In the belief-based AI conditions, the system was presented as autonomously selecting the strategy and trajectory, while participants initiated the trial by pressing the space bar. Thus, all conditions involved a single keypress at trial onset, ensuring comparable motor requirements across conditions and allowing differences in behavioural and neural responses to be more confidently attributed to agency-related manipulations rather than to variations in motor output.

Immediately after the keypress, the car moved towards the selected target. In all conditions, the car’s movement lasted 300 ms. Upon reaching the destination, an auditory tone signalled the trial outcome. Auditory feedback consisted of two 100-ms pure tones delivered at 80 dB SPL: a high-pitched tone (1000 Hz) and a low-pitched tone (500 Hz). In 80% of trials, the tone matched the expected outcome based on the selected strategy (match trials), whereas in the remaining 20% of trials, an unexpected tone was presented to elicit auditory prediction-error responses (mismatch trials).

To maintain attention and enhance oddball-related ERP components, a subset of trials served as catch trials, in which tone amplitude was reduced by 50%. Participants were instructed to respond as quickly as possible to these catch trials using the same key used for trial initiation, and performance feedback was provided at the end of each block.

In 20% of trials, 1200 ms after tone offset, participants additionally rated their feeling of control by answering the question: “How much did you feel in control over the trial?” Responses were given on a Likert scale ranging from 1 (no control) to 9 (complete control). These explicit ratings were included to verify task engagement and to confirm that the experimental manipulations produced the expected behavioural effects of automation and explanatory support observed in previous work using this paradigm (Houdoyer et al., under review). The proportion of rating trials was intentionally limited to minimise disruption of task flow and preserve the integrity of the electrophysiological recordings. Accordingly, these ratings were not intended for trial-level correlations with EEG measures, but served as a manipulation check and behavioural validation of the experimental design. A 700-ms blank screen preceded the next trial.

The three experiments shared this common auditory-feedback structure and trial sequence, differing only in the source of decision control and the type of explanation provided by the purported AI system. The specific manipulations implemented in each experiment are detailed below.

#### Belief-Based AI Assistance Context

Across all belief-based AI conditions, target selection and vehicle movement were presented as being controlled by an AI-based driving assistance system. The task was designed to simulate a high-level automated driving situation in which users did not continuously control the vehicle, but instead configured the assistance system, monitored its behaviour, and validated the final outcome.

Before the automated blocks, participants were informed that they would interact with an artificial intelligence system capable of selecting an appropriate strategy and trajectory on each trial. The system was described as a decision-tree AI designed to optimise three competing parameters: safety, reward, and driving time. To reinforce this interaction context, participants configured the purported AI system before the automated task by indicating the level of risk they were willing to accept. This risk preference was expressed as the probability of encountering an obstacle and ranged from 15% to 50%. Participants were told that lower risk settings favoured safer but slower and less rewarding trajectories, whereas higher risk settings favoured faster and more rewarding but riskier trajectories. They were also informed that this preference would be integrated with trial-specific task parameters when the AI computed the vehicle’s strategy and trajectory.

In reality, the vehicle’s behaviour was pre-programmed to ensure experimental control and standardisation across participants. Strategy and trajectory selections followed predefined, counterbalanced schedules rather than being dynamically adapted to participants’ responses or risk preferences. Thus, participants’ settings did not modify the system’s behaviour online, but served to support the belief that the AI’s decisions were shaped by both task constraints and user-defined preferences. The paradigm should therefore be understood as a controlled, belief-based simulation of conditional automation, conceptually inspired by SAE Level 3 automated driving, rather than as a technically implemented SAE Level 3 system.

#### Experiment 1: Effect of automation on the sense of agency (Motor/no-AI vs. belief-based AI)

Experiment 1 examined how automation influences the SoA by contrasting self-determined actions with actions attributed to a belief-based AI assistance system (see Figures 1A and 1B). Participants completed two within-subject conditions, Motor/no-AI and belief-based AI, presented in alternating blocks, with block order counterbalanced across participants.

In the Motor/no-AI condition, participants selected the strategy themselves and initiated the trial by pressing the arrow key corresponding to the spatial location of the chosen target. In the belief-based AI condition, target selection was presented as being controlled by an AI-based driving assistance system. Participants were informed that the system selected the strategy by optimising task-relevant parameters, including safety, reward, and driving time, while taking into account the risk preference they had configured before the automated block. In reality, the AI’s choices followed a predefined and counterbalanced schedule to ensure experimental control and standardisation across participants.

Participants no longer determined the strategy directly in the belief-based AI condition. Their role was to monitor the AI-driven decision process, initiate the trial by pressing the space bar, respond to occasional obstacle events when required, and validate the final outcome. Thus, trial initiation was required in both conditions, allowing the effect of decision autonomy to be isolated while maintaining comparable motor involvement across conditions.

#### Experiment 2: Effect of explanation transparency in automated decisions

Experiment 2 tested whether distal-level explanations, corresponding to the rationale underlying the selected strategy, modulated the SoA during belief-based AI decision-making (see Figures 1B–1D). All decisions were presented as being made by the AI-based assistance system. Participants completed two within-subject conditions, belief-based AI without explanation and belief-based AI with distal explanation, presented in alternating blocks, with block order counterbalanced across participants.

Before the automated blocks, participants were informed that the AI-based assistance system selected its behaviour by balancing safety, reward, and driving time while taking into account their previously configured risk preference. Explanations were framed as concise, task-specific statements indicating that the AI’s choice was advantageous with respect to these decision-relevant parameters. Because the task involved rapid events and limited processing time, explanations were designed to be immediately interpretable. They therefore combined brief feature-related textual statements, such as “Gain” and “Security”, with visual cues indicating the AI’s upcoming decision.

In the belief-based AI without explanation condition, both targets were displayed with green checkmarks and an uninformative placeholder text (“Null/Inconnu”) for 600 ms. This condition served as a visual control for the presence of explanatory cues. The spatial arrangement and display duration of the checkmarks and text were matched to those of the explanation condition, but the placeholder did not provide meaningful information about the AI’s decision.

In the belief-based AI with distal explanation condition, the target selected by the AI was highlighted in yellow and accompanied by brief feature-related statements indicating the task-relevant reasons for the selected strategy, such as “Gain” and “Security”, displayed with green checkmarks. This information was presented for 600 ms before trial initiation. The target highlight provided direct visual information about the AI’s goal-level decision, namely the selected strategy, while the textual statements specified why this strategy was advantageous with respect to the task parameters.

In both conditions, the car icon then appeared at the centre of the screen, indicating that participants could initiate the trial by pressing the space bar. Participants had no control over the selected strategy in either condition, ensuring equivalent motor engagement while isolating the effect of meaningful explanatory content.

#### Experiment 3: Effect of combining proximal and distal explanations

Experiment 3 investigated whether adding proximal-level information, corresponding to the selected trajectory, enhanced the effect of distal-level explanations, corresponding to the selected strategy, on the SoA during belief-based AI decision-making (see Figures 1B–1E). The general trial structure, timing, auditory feedback, and motor requirements were identical to those used in Experiments 1 and 2, except that the visual layout and explanatory cues were adapted to introduce proximal ambiguity.

Each trial displayed four targets, two per strategy, positioned in the four corners of the screen. This layout allowed two possible trajectories for each strategy and therefore made it possible to dissociate strategy-level and trajectory-level information. Participants completed three within-subject conditions, with condition order counterbalanced across participants: belief-based AI without explanation, belief-based AI with distal explanation, and belief-based AI with proximal + distal explanation.

As in Experiment 2, explanations consisted of concise, task-specific statements indicating that the AI’s choice was advantageous with respect to the decision-relevant parameters, namely reward, safety, and driving time. These statements were displayed as “Gain” and “Security” accompanied by green checkmarks, and were combined with visual highlights around the relevant target or targets to indicate the intentional level addressed by the explanation.

In the belief-based AI without explanation condition, the four targets were displayed without informative highlighting, and a central placeholder text served as a visual control for the presence of explanatory information.

In the belief-based AI with distal explanation condition, the two targets corresponding to the AI-selected strategy were highlighted in yellow. This provided information about the AI’s broader goal-level decision, namely the selected strategy, while leaving the specific trajectory ambiguous. The target highlights were accompanied by the same brief feature-related statements, such as “Gain” and “Security”, indicating why the selected strategy was advantageous according to the task parameters.

In the belief-based AI with proximal + distal explanation condition, only one of the two targets corresponding to the AI-selected strategy was highlighted. This provided both distal information about the selected strategy and proximal information about the specific trajectory selected by the AI. The textual explanation was identical in format to that used in the distal-explanation condition, allowing the two explanation conditions to differ primarily in the precision of the visual information provided about the AI’s upcoming decision.

After the 600-ms presentation of the visual and textual cues, the car appeared at the centre of the screen, prompting participants to initiate the trial by pressing the space bar. As in the previous belief-based AI conditions, the AI was presented as autonomously determining the selected strategy and trajectory, while participants only initiated the movement. The car then moved toward the selected target. All auditory feedback, timing parameters, and core visual elements were kept constant across experiments to ensure methodological comparability.

### 2.4 Experimental Factors

#### Tone Predictability

Across all three experiments, Tone Predictability was included as a common within-subject factor. This factor contrasted match trials, in which the auditory feedback was consistent with the outcome expected from the selected strategy, with mismatch trials, in which the auditory feedback violated this expectation. Tone Predictability was included to assess whether auditory prediction errors modulated Judgment of Control ratings. Thus, this factor captured the extent to which participants’ perceived control was sensitive to violations of the expected sensory consequences of the selected or AI-selected action.

#### Automation

In Experiment 1, the degree of AI-based assistance varied across conditions. In the Motor/no-AI condition, participants selected the strategy themselves and initiated the vehicle movement using the arrow key corresponding to the spatial location of the chosen target. In the belief-based AI condition, target selection was presented as being controlled by the AI-based driving assistance system. Participants no longer determined the strategy directly. Instead, their role was to monitor the purported AI-based decision process, initiate the trial using the space bar, respond to occasional obstacles or catch events when required, and validate the final outcome. Importantly, all conditions involved a single keypress at trial onset, ensuring that differences in behavioural and neural responses could be more confidently attributed to agency-related manipulations rather than to differences in motor output. This manipulation, therefore, contrasted self-determined actions with actions attributed to the belief-based AI system, allowing us to test whether reduced decision autonomy altered Judgment of Control.

#### AI Explainability in Experiment 2

In Experiment 2, the experiment-specific factor was AI Explainability. All trials were presented as being controlled by the belief-based AI system, and the manipulation contrasted trials without explanation with trials accompanied by a distal explanation. During the automated task, the purported AI-based assistance system was presented as selecting its behaviour by optimising task-relevant parameters, namely safety, reward, and driving time, while taking into account the participant’s previously configured risk preference.

Consistent with feature-based approaches to explainable AI, explanations were operationalised as concise, human-interpretable information about the parameters presented as guiding the system’s decision. In the belief-based AI with distal explanation condition, explanations consisted of brief task-specific statements indicating that the AI’s choice was advantageous with respect to the decision-relevant parameters. These statements were displayed as “Gain” and “Security”, accompanied by green checkmarks. Because the task involved rapid events and limited processing time, the explanations were designed to be immediately interpretable. They were therefore combined with a visual cue indicating the AI’s upcoming goal-level decision: the target selected by the AI was highlighted in yellow. This distal explanation provided information about the selected strategy, that is, the broader goal toward which the AI’s decision was directed.

In the belief-based AI without explanation condition, no meaningful information about the AI’s upcoming decision was provided. Instead, non-informative visual cues and placeholder text were presented for the same duration and in a matched visual format, serving as a control for the mere presence of additional cues and text. Thus, the AI Explainability factor in Experiment 2 tested whether strategy-level information about the AI’s decision modulated Judgment of Control during belief-based AI decision-making.

#### AI Explainability in Experiment 3

In Experiment 3, AI Explainability again served as the experiment-specific factor, but comprised three levels: belief-based AI without explanation, belief-based AI with distal explanation, and belief-based AI with combined proximal + distal explanation. The four-target layout introduced proximal ambiguity by allowing two possible trajectories for each strategy. This made it possible to dissociate information about the AI’s broader goal-level decision from information about its immediate action-level decision.

In the belief-based AI without explanation condition, no meaningful information about the AI’s upcoming decision was provided; only non-informative visual-control cues were displayed. In the belief-based AI with distal explanation condition, the two targets corresponding to the AI-selected strategy were highlighted in yellow. This provided information about the AI’s broader goal-level decision, namely the selected strategy, while leaving the specific trajectory ambiguous. In the belief-based AI with combined proximal + distal explanation condition, only the specific target selected by the AI was highlighted. This provided information both about the selected strategy and about the selected trajectory, corresponding to the AI’s immediate action-level decision.

In both explanation conditions, the visual cue was accompanied by the same concise feature-related statements, such as “Gain” and “Security”, displayed with green checkmarks. Thus, the two explanation conditions differed primarily in the precision of the visual information provided about the AI’s upcoming decision, while preserving the same textual explanation format. These explanations were prospective, pre-action cues presented before movement onset, rather than retrospective justifications delivered after the action had been completed. The manipulation therefore targeted participants’ anticipation and monitoring of the AI’s upcoming behaviour, rather than providing a full mechanistic account of the algorithm’s internal computations. This design allowed us to examine whether increasing the intentional specificity of information about the purported AI’s decision modulated Judgment of Control.

### 2.5 Group Assignment and Procedure

Each experiment was conducted on an independent cohort, with no overlap between participants. Participants were randomly assigned to Experiment 1, Experiment 2, or Experiment 3. In Experiments 1 and 2, participants completed eight blocks of 50 trials per condition, resulting in 400 trials per condition. Each block included 40 match trials, in which the auditory feedback corresponded to the expected outcome, and 10 mismatch trials, in which the auditory feedback was unexpected. Thus, mismatch trials represented 20% of trials in each block. Experimental blocks alternated between conditions, for example Motor/no-AI → belief-based AI → Motor/no-AI → belief-based AI in Experiment 1, with the starting condition counterbalanced across participants.

Experiment 3 followed a different block structure. The experiment consisted of eight blocks, with two blocks administered for each map configuration. Each block was subdivided into three successive sub-blocks corresponding to the three experimental conditions: belief-based AI without explanation, belief-based AI with distal explanation, and belief-based AI with combined proximal + distal explanation. This resulted in alternating condition sequences throughout the session. The order of sub-blocks was counterbalanced across participants to control for order effects. Each sub-block contained 34 trials, comprising 26 match trials and 8 mismatch trials.

In the belief-based AI conditions, 10% of trials were designated as obstacle trials. These trials were randomly interspersed throughout the session. Obstacle trials also served as catch trials, in which the auditory tone amplitude was reduced by 50%. Participants were instructed to respond to these trials as quickly as possible. In the Motor/no-AI condition, obstacle occurrence depended on the strategy selected by the participant and therefore followed the obstacle probability associated with the chosen strategy rather than a fixed predefined rate.

In addition, 20% of all trials included a sense-of-agency (SoA) rating, presented 1200 ms after auditory feedback. Participants rated their feeling of control over the trial using the scale described above. The mapping between auditory pitch and expected versus unexpected outcomes was counterbalanced across participants to prevent frequency-specific confounds. Before the main task, participants completed a short practice phase consisting of 10 trials per condition. This practice phase familiarised participants with the task structure, response mappings, and feedback contingencies. Short breaks were provided between blocks to minimise fatigue and maintain high-quality EEG recordings throughout the session.

### 2.6 EEG Acquisition and Preprocessing

EEG signals were recorded using a Neuroelectrics Enobio 20 system with a custom 17-channel montage (Fp1, Fpz, Fp2, F7, F3, Fz, F4, F8, C3, Cz, C4, T7, P7, Pz, P8, T8, Oz – see figure 3.E.). Electrode positions were digitized in three-dimensional Cartesian space to ensure accurate spatial localization. Data were sampled at 500 Hz (effective rate 500.01 Hz) via USB using NIC software (version 2.1.3.11, macOS) and stored in.easy format. A 50 Hz online infinite impulse response (IIR) notch filter was applied during acquisition to suppress mains interference. No additional online filtering, re-referencing, or EOG correction was applied. EEG signals were acquired using the system’s standard hardware reference configuration based on common mode sense and driven right leg (CMS/DRL) electrodes. EEG amplitudes were recorded in nanovolts (nV). A three-axis accelerometer (100 Hz) was recorded concurrently but was not analyzed further.

Preprocessing was performed in MNE-Python (1.11.0) using a standardised, semi-automated pipeline. Continuous EEG data were first visually inspected to identify gross artefacts before automated cleaning. Noisy channels were identified using a composite detection approach combining statistical and correlation-based metrics, supplemented by PyPREP’s automated diagnostics. Channels that consistently deviated from the global signal pattern were excluded prior to ICA (0.68 ±1.06 bad channels per participant in Experiment 1, 1.39 ±1.50 in Experiment 2, and 1.39 ±1.50 in Experiment 3) and later interpolated. This procedure ensured robust artefact detection while minimising false positives.

Independent component analysis (ICA) was performed using FastICA, retaining approximately 99% of the total signal variance. Components reflecting ocular activity (blinks and eye movements) and muscle artefacts were identified and removed before reconstructing the cleaned signal. On average, 1.35 ± 0.67 components were removed in Experiment 1, 1.67 ± 0.82 in Experiment 2, and 2.11 ±0.88 in Experiment 3.

Following ICA, previously excluded channels were reintroduced and interpolated using spherical splines. To attenuate residual mains contamination and its first harmonic not fully suppressed by the online 50 Hz notch applied during acquisition, the data were filtered offline using 50 and 100 Hz notch filters. The data were then band-pass filtered from 0.1 to 80 Hz using a zero-phase FIR filter implemented in MNE-Python, thereby reducing slow drifts and high-frequency noise while preserving phase information. EEG epochs were extracted from - 100 to +600 ms relative to auditory feedback onset and baseline-corrected using the-100 to 0 ms interval.

Trials exceeding ±150 µV were automatically rejected. For each participant, grand-average ERPs were computed separately for all experimental conditions within each study (e.g., Motor/no-AI-Match, AI without explanation-Match, Motor/no-AI-Mismatch, AI without explanation-Mismatch, etc.). Only regular, successfully completed trials were retained for ERP averaging. Regular trials were defined as trials without obstacle events and without catch-tone detection demands, whereas successful trials were defined as trials in which participants produced a single appropriate response, with no additional keypresses suggesting overshooting or an inappropriate response strategy. Catch and obstacle trials were excluded from all ERP analyses to avoid contaminating the averaged waveforms with attentional or motor responses unrelated to the main experimental manipulation.

### 2.7 Data Analysis

#### 2.7.1 Behavioural Data Analysis

Behavioural analyses were conducted in R (version 4.5.2). Linear mixed-effects models were fitted using the `lme4` package, with degrees of freedom and p-values approximated using Satterthwaite’s method. Post hoc pairwise comparisons were conducted using Tukey-adjusted t-tests to correct for multiple comparisons.

Only standard trials were included in the behavioural analyses. Catch trials and obstacle trials were excluded, as they were not treated as trials of interest. Missing values were not imputed. Trials with missing Judgment of Control ratings were excluded only from the corresponding behavioural analysis, and models were fitted on all available valid observations. Judgment of Control (JoC) measures were Z-standardized to account for inter-individual differences in response scaling. Z-scores were computed as:

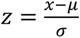

*μ* and *σ* denote the participant-specific mean and standard deviation across valid Judgment of Control trials, respectively. Trials with absolute z-scores exceeding 2 ((|z| > 2)) were treated as outliers and excluded from further analyses. This outlier-removal procedure reduced the Judgment of Control dataset from 4,720 to 4,536 trials in Experiment 1, from 4,738 to 4,630 trials in Experiment 2, and from 6,426 to 6,175 trials in Experiment 3.

Linear mixed-effects models were used to analyse trial-level Judgment of Control ratings. This modelling approach was chosen because the data had a repeated-measures structure, with multiple trials nested within each participant. Mixed-effects models allowed trial-level observations to be retained while accounting for the non-independence of observations from the same participant through participant-level random effects.

All models included random intercepts for participants, allowing each participant to have their own baseline level of Judgment of Control. Random slopes for within-subject predictors were included whenever supported by model convergence and retained unless they led to convergence failures or singular fits. Although models were fitted to trial-level data, individual trials were therefore not treated as independent observations; the dependency among repeated observations from the same participant was explicitly modelled through the random-effects structure.

For Experiment 1, fixed effects included Automation, Tone Predictability, and their interaction. Automation comprised two levels: Motor/no-AI and belief-based AI. For Experiment 2, fixed effects included AI Explainability, Tone Predictability, and their interaction. AI Explainability comprised two levels: belief-based AI without explanation and belief-based AI with distal explanation. For Experiment 3, fixed effects included AI Explainability, Tone Predictability, and their interaction. AI Explainability comprised three levels: belief-based AI without explanation, belief-based AI with distal explanation, and belief-based AI with combined proximal + distal explanation.

For all models, significant interactions were followed up using Tukey-adjusted pairwise comparisons. Partial eta-squared values for mixed-effects models were computed using the **effectsize** package from the ANOVA tables obtained from the fitted models.

#### 2.7.2 ERP Data Analysis

ERP analyses focused on the auditory-evoked P1, N1, and P2 components, as these fronto-central responses are known to be modulated by sensory attenuation and predictive mechanisms underlying the SoA (Kühn et al., 2011; Timm et al., 2014; Bednark et al., 2015). For each participant, ERPs were computed separately for each experimental condition within the corresponding study (e.g., Motor/no-AI-Match, Motor/no-AI-Mismatch, belief-based AI without explanation-Match, and belief-based AI without explanation-Mismatch in Experiment 1). EEG epochs were time-locked to tone onset (−100 to 600 ms) and baseline-corrected using the −100 to 0 ms interval.

Grand-average waveforms were then computed across participants for visualization. Component peaks and latencies were extracted from a predefined fronto-central region of interest (ROI; F3, Fz, F4, C3, Cz, C4, Pz), where auditory P1–N1–P2 components typically reach their maximal amplitude. For each participant, condition, and electrode, peak amplitudes were measured within predefined time windows (P1: 30-70 ms; N1: 80-120 ms; P2: 150-250 ms). The P1 and P2 peaks were defined as the maximal positive deflections within their respective windows, whereas the N1 peak was defined as the most negative deflection. Corresponding peak latencies (ms) were extracted from the time indices of these extrema. To characterise the dynamics of early auditory processing associated with the SoA, both individual component measures (P1, N1, P2) and composite measures were computed. The latter included P1–N1 and N1–P2 peak-to-peak amplitudes, providing an integrated quantification of early sensory encoding and feedback-related responses associated with sensory attenuation mechanisms.

For each dependent variable (peak amplitude, latency, and composite measures), repeated-measures ANOVAs were conducted. In Experiment 1, analyses included Action (Motor/no-AI vs belief-based AI), Tone Predictability (Match vs Mismatch) within-subject factors. In Experiment 2, the Action factor was replaced by AI Explainability, defined as belief-based AI without explanation versus belief-based AI with distal explanation, while Tone Predictability remained within-subject factors. In Experiment 3, AI Explainability included three levels (belief-based AI without explanation, belief-based AI with distal explanation, and belief-based AI with combined proximal + distal explanation), again crossed with Tone Predictability as within-subject factors.

To further examine neural responses to auditory prediction errors, the Mismatch Negativity (MMN) component was computed as the difference between mismatch and match ERPs (Mismatch - Match), separately for each condition and participant. This subtraction isolates automatic prediction-error processing while minimizing contamination from exogenous auditory responses.

Analyses were performed using the same 17-channel montage and preprocessing pipeline described above. Following standard MMN conventions, analyses focused on the 100-250 ms post-stimulus interval, corresponding to the canonical latency range of auditory prediction-error responses (Näätänen et al., 2007). Within this window, two parameters were extracted for each participant, condition, and electrode within a restricted fronto-central ROI (F3, Fz, F4, C3, Cz), where the MMN typically reaches its maximal amplitude: (i) peak amplitude (µV), defined as the most negative deflection of the difference waveform; and (ii) peak latency (ms), corresponding to the time point of this maximal deflection.

For MMN measures, repeated-measures ANOVAs included Action (e.g., Motor/no-AI vs belief-based AI in Experiment 1) or AI Explainability (belief-based AI with vs without explanation in Experiment 2; belief-based AI distal, belief-based AI proximal + distal, and belief-based AI without explanation in Experiment 3) as within-subject factors. Post-hoc pairwise comparisons were corrected for multiple testing using the Benjamini-Hochberg FDR procedure, and Hedges’ g was reported as an effect-size estimate.

## 3. Results

### 3.1 Self-report Measure: Feeling of Control (FoC)

Participants’sense of agency was assessed on 20% of trials using a 9-point scale rating their perceived control over the car’s movement.

#### 3.1.1 Effect of Automation on Perceived Control: Experiment 1

Automation produced a robust reduction in perceived control (F(1, 31.4) = 16.25, p = 0.00033, η²ₚ = 0.34) (see fig.2.A). Neither the main effect of Tone Predictability, F(1, 431.4) = 0.0004, p = 0.98, nor the Action × Tone Predictability interaction, F(1, 4476.5) = 0.42, p = 0.51, reached significance.

**Figure 2.**
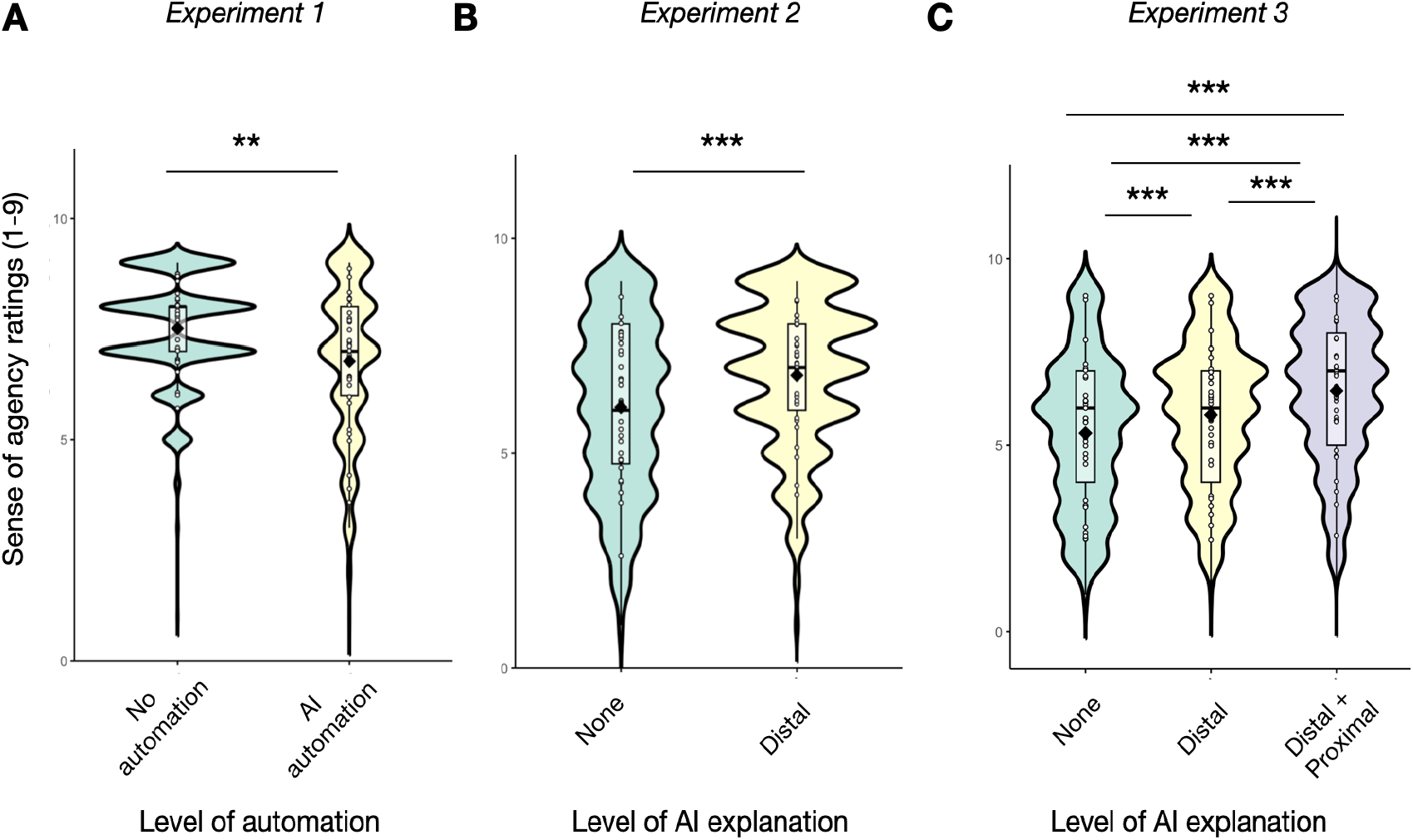
Impact of automation and AI explanations on sense of agency (SoA) ratings. (A) Effect of automation level (no automation vs. automation) on explicit SoA ratings. (B) Effect of distal explanation (No-Explanation vs. distal) on explicit SoA ratings. (C) Effect of AI explanations combination (No-Explanation, Distal, and Combined) on explicit SoA ratings. Overall, automation significantly reduced perceived agency, whereas system explainability improved ratings, particularly when Proximal and Distal rationales were combined; N.B.: * = p < 0.05; ** = p < 0.01; *** = p < 0.001; **** = p < 0.0001.

**Figure 3.**
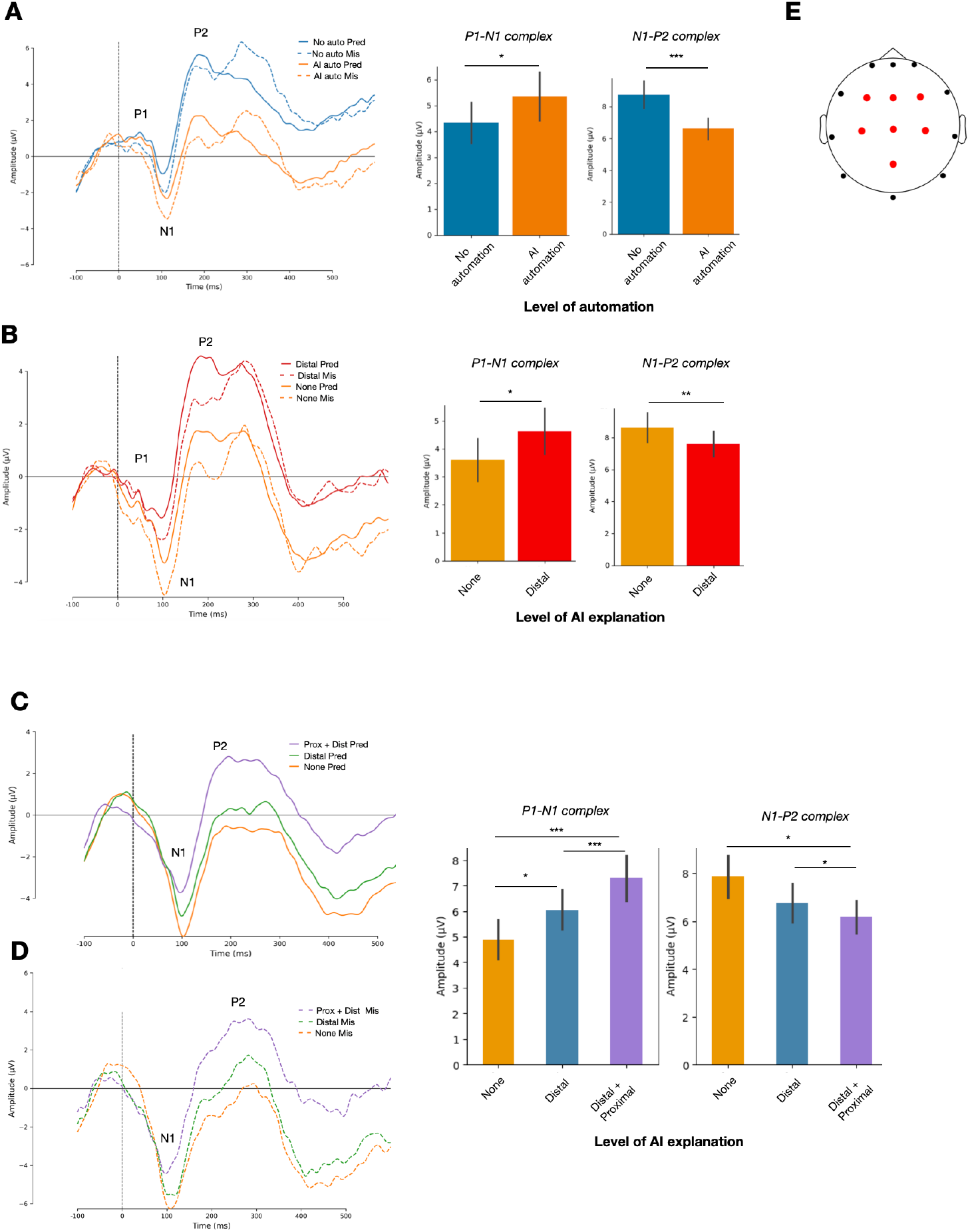
Event-related potentials as a function of automation and AI explanation. **A**, Grand-average event-related potentials (ERPs) in the no-automation and AI-automation conditions of Experiment 1, averaged across the frontocentral region of interest (ROI). Solid traces indicate predicted tones (Pred; regular tones), whereas dashed traces indicate mispredicted tones (Mis; oddball tones). Bar plots show the peak-to-peak amplitudes of the P1–N1 and N1–P2 complexes for each automation condition. **B**, Grand-average ERPs in the no-explanation and distal-explanation conditions of Experiment 2, shown separately for Pred and Mis trials. Bar plots show P1–N1 and N1–P2 peak-to-peak amplitudes for each explanation condition. c,d, Grand-average ERPs for Pred (**C**) and Mis (**D**) trials in Experiment 3, displayed for the no-explanation, distal-explanation, and combined distal-plus-proximal-explanation conditions. The corresponding bar plots show P1–N1 and N1–P2 peak-to-peak amplitudes across explanation conditions. **E**, Electrode montage showing the frontocentral ROI used for ERP quantification (F3, Fz, F4, C3, Cz, C4 and Pz; red). Time zero denotes tone onset. N.B.: * = p < 0.05; ** = p < 0.01; *** = p < 0.001; **** = p < 0.0001.

Tukey-corrected post-hoc comparisons confirmed that FoC ratings were significantly higher in the Motor/no-AI condition (M = 7.52, SE = 0.15, 95% CI [7.22, 7.82]) than in the belief-based AI condition (M = 6.80, SE = 0.26, 95% CI [6.27, 7.33]; β = 0.721, t(31.8) = 3.57, p = 0.0012). These results demonstrate that delegating control to the belief-based AI substantially diminished participants’ explicit sense of control (Figure 2A). To determine whether this reduction could be mitigated, we next examined the impact of explanatory information, first through distal (goal-level) explanations and subsequently through combined distal + proximal (goal-and path-level) explanations.

#### 3.1.2 Effect of Distal Explanation Type on Perceived Control: Experiment 2

Providing distal explanations (see fig.2.B) had a robust positive effect on participants’ perceived control (F(1, 31.5) = 15.83, p = 0.00038, η²ₚ = 0.32). Consistent with Experiment 1, the main effect of Tone Predictability was not significant, F(1, 1990.5) = 2.29, p = 0.13, nor was the AI Explainability × Tone Predictability interaction, F(1, 4560.5) = 1.51, p = 0.22.

Tukey-corrected comparisons showed that FoC ratings were significantly lower in the No-Explanation condition (M = 6.08, SE = 0.28, 95% CI [5.50, 6.65]) than when the belief-based AI disclosed its distal, goal-level rationale (M = 6.81, SE = 0.22, 95% CI [6.36, 7.26]). This improvement was further supported by the mixed-effects model (β =-0.735, t(32.5) = −3.91, p = 0.0006). Together, these results indicate that making the belief-based AI’s high-level objectives explicit significantly enhances participants’ perceived control relative to an opaque system.

#### 3.1.3 Effect of combined Distal and Proximal Explanation on Perceived Control: Experiment

In Experiment 3, explanation type (see fig.2.C) also had a strong effect on FoC ratings (F(2, 37.8) = 18.61, p = 1.57×10⁻⁷, η²ₚ = 0.56). As in Experiments 1 and 2, the main effect of Tone Predictability was not significant, F(2, 667.3) = 0.46, p = 0.50, nor was the AI Explainability × Tone Predictability interaction, F(2, 6074.6) = 1.14, p = 0.32.

Participants reported substantially lower perceived control in the No-Explanation condition (M = 5.34, SE = 0.31, 95% CI [4.72, 5.96]) compared with either explanatory format. Distal explanations alone significantly increased perceived control (M = 5.84, SE = 0.28, 95% CI [5.27, 6.41]; β = −0.500, t(37.1) = −4.40, p <.0001). The highest FoC ratings were observed when proximal and distal information were combined (M = 6.42, SE = 0.28, 95% CI [5.85, 6.98]; β = −1.077, t(35.8) = −5.82, p <.0001).

Crucially, Tukey-corrected comparisons showed that combined explanations significantly improved perceived control relative to distal explanations alone (β = −0.577, t(37.6) = −5.72, p <.0001), demonstrating an additive benefit of integrating both goal-level and path-level cues. Across Experiments 2 and 3, these results converge to show that while explanations reliably enhance participants’ SoA, dual-level intention sharing yields the strongest and most consistent improvement.

### 3.2 ERP

#### 3.2.1 Effect of Automation on Perceived Control: Experiment 1

Repeated-measures ANOVAs were conducted with Action (Motor/no-AI, belief-based AI) and Tone Predictability (Match, Mismatch) as within-subject factors, together with their interaction. Analyses focused on P1–N1 and N1–P2 peak-to-peak amplitudes, MMN amplitude, and the latencies of the P1, N1, and P2 peaks, as well as the MMN component (see Supplementary Results, “Latency Analyses,” Section 1). The principal findings are summarised below, with Figure 3.A illustrating the peak-to-peak complexes and Figure 4.A the MMN component.

**Figure 4.**
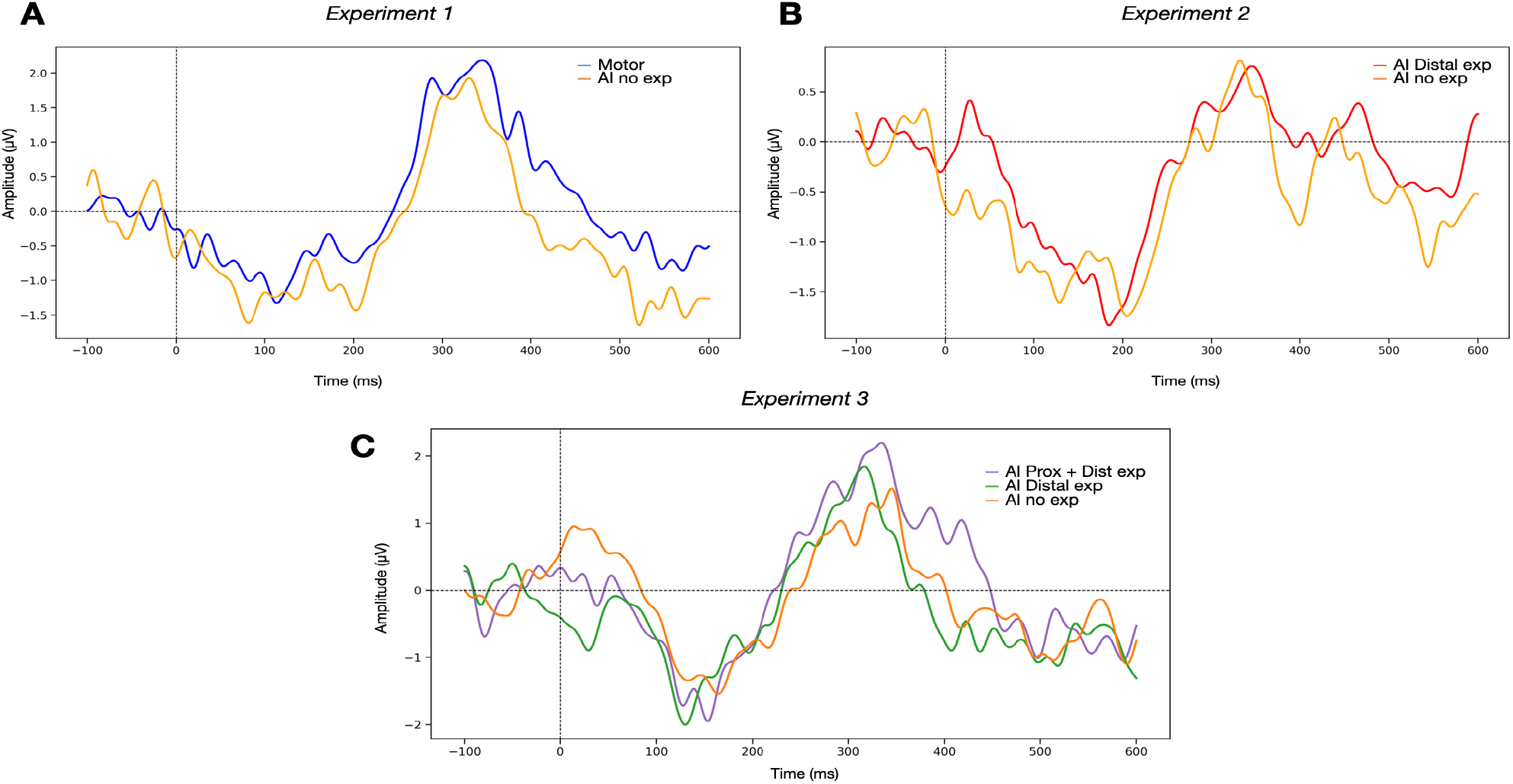
Mismatch Negativity (MMN) across experimental conditions. (A) Grand-average MMN waveforms for the Motor and AI no-explanation conditions (Experiment 1). (B) Grand-average MMN waveforms for the AI distal explanation and no-explanation conditions (Experiment 2). (C) Grand-average MMN waveforms for the AI no-explanation, distal explanation, and combined proximal + distal explanation conditions (Experiment 3). Waveforms are time-locked to stimulus onset (0 ms; vertical dashed line), with amplitude expressed in µV. The horizontal dashed line indicates zero amplitude. Overall, MMN responses varied across conditions, with differences observed between explanation levels and relative to the motor condition.

##### P1–N1 Peak-to-Peak Amplitude

A significant main effect of Action was observed, F(1,19) = 4.97, p = 0.0374, η²ₚ = 0.023, with overall larger amplitudes in the belief-based AI-assisted condition than in the Motor/no-AI condition (post hoc: t(19) = 2.26, p = 0.0374, Hedges’ g = 0.305). A significant main effect of Tone Predictability was also found, F(1,19) = 6.91, p = 0.0161, η²ₚ = 0.018, with mismatch tones producing larger P1–N1 amplitudes than match tones (post hoc tests: t(19) = –3.74, p = 0.0016, Hedges’ g = –0.273). The Action × Tone Predictability interaction did not reach significance, F(1,19) = 0.053, p = 0.82.

##### N1–P2 Peak-to-Peak Amplitude

Analysis of the N1–P2 complex revealed a significant main effect of Action, F(1,19) = 25.32, p = 6.4 × 10⁻⁵, η²ₚ = 0.100, with markedly smaller amplitudes in the belief-based AI-assisted condition than in the self-initiated condition (post hoc: t(19) = –4.16, p <.001, Hedges’ g = – 0.406). No significant main effect of Tone Predictability was observed, F(1,19) = 0.48, p =.50, although mismatch tones elicited numerically larger responses. The Action × Tone Predictability interaction did not reach significance, F(1,19) = 1.11, p =.30.

##### MMN peak

To assess whether automation influenced pre-attentive deviance processing, mismatch negativity (MMN) peak amplitude was compared between the Motor/no-AI and belief-based AI-assisted conditions (see Figure 4A). This analysis revealed no significant effect of Condition on MMN peak amplitude, F(1,19) = 0.30, p =.592, η²ₚ = 0.004. Accordingly, the present data do not provide evidence that automation altered this index of pre-attentive auditory deviance processing in Experiment 1.

#### 3.2.2 Effect of Distal Explanation Type on Perceived Control: Experiment 2

As in Experiment 1, repeated-measures ANOVAs were conducted with AI Explainability (no explanation, distal explanation) and Tone Predictability (Match, Mismatch) as within-subject factors, along with their interaction. Analyses focused on P1–N1 and N1–P2 peak-to-peak amplitudes (Figure 3B), MMN amplitude (Figure 4B), and the latencies of the P1, N1, and P2 peaks, as well as of the MMN component (see Supplementary Results, “Latency Analyses,” Section 2.)

##### P1–N1 Peak-to-Peak Amplitude

The analysis revealed a significant main effect of AI Explainability (F(1,17) = 5.10, p = 0.037), indicating reduced amplitudes when distal explanations were provided (t(17) = −2.26, p = 0.0374, Hedges’ g = −0.26). A significant main effect of Tone Predictability (F(1,17) = 13.97, p = 0.0016, η²ₚ = 0.035) was also observed, with mismatch tones eliciting larger P1–N1 amplitudes than match tones (t(17) = −3.74, p = 0.0016, Hedges’ g =-0.37). The AI Explainability × Tone Predictability interaction was not significant (F(1,17) = 0.13, p = 0.72).

##### N1–P2 Peak-to-Peak Amplitude

Analysis of the N1–P2 complex revealed a main effect of AI Explainability (F(1,17) = 12.06, p = 0.0029, η²ₚ = 0.0206), with significantly smaller amplitudes in the no-explanation condition than in the distal-explanation condition (t(17) = −3.47, p = 0.0029, Hedges’ g = −0.28). Neither the main effect of Tone Predictability (F(1,17) = 1.23, p = 0.28) nor the AI Explainability × Tone Predictability interaction (F(1,17) = 0.95, p = 0.34) reached significance (see Figure 3.B).

##### MMN peak

To examine whether distal explanations influenced pre-attentive deviance detection, MMN peak amplitude (see Figure 4.B) was compared between the No-explanation and Distal-explanation conditions. No significant difference was found for MMN peak amplitude (F(1,17) = 0.17, p = 0.682, η²ₚ = 0.004). Thus, unlike earlier ERP components, the MMN response remained stable across conditions, suggesting that pre-attentive deviance detection processes were unaffected by whether the belief-based AI disclosed its distal intentions.

#### 3.2.3 Effect of combined Distal and Proximal Explanation on Perceived Control: Experiment 3

Consistent with the preceding experiments, repeated-measures ANOVAs were conducted with AI Explainability (no explanation, distal explanation, combined explanation), Tone Predictability (Match, Mismatch), and their interaction as within-subject effects. Analyses focused on P1–N1 and N1–P2 peak-to-peak amplitudes (Figure 3.C-D), MMN amplitude (Figure 4.C), and the latencies of the P1, N1, and P2 peaks, as well as of the MMN component (see Supplementary Results, “Latency Analyses,” Section 3).

##### P1–N1 Peak-to-Peak Amplitude

The analysis revealed a strong influence of both AI Explainability and Tone Predictability on the P1–N1 complex. Critically, the level of explanation provided by the belief-based AI exerted a robust effect on early auditory processing, as indicated by a strong main effect of AI Explainability (F(2,36) = 24.30, p = 4.1 × 10⁻⁶, η²ₚ = 0.13). A significant main effect of Tone Predictability was also found (F(1,18) = 16.79, p = 0.00068, η²ₚ = 0.040). The AI Explainability × Tone Predictability interaction did not reach statistical significance (F(2,36) = 2.52, p = 0.10), indicating that explanation-related differences were consistent across tone types.

Post-hoc tests clarified the directionality of these effects. A clear graded pattern of sensory attenuation was also observed across explanation levels: amplitudes were significantly smaller in the distal-explanation condition compared to the no-explanation condition (t(18) = −2.77, p = 0.0126, Hedges’g = −0.25). This attenuation was even more pronounced in the combined proximal + distal explanation condition, which showed markedly reduced amplitudes relative to both the no-explanation condition (t(18) = −6.19, p = 7.6 × 10⁻⁶, Hedges’g = −0.84) and the distal-only condition (t(18) = 3.95, p = 0.00093, Hedges’g = 0.60). Regarding Tone Predictability, Mismatch tones (Fig 3.D) elicited significantly larger P1–N1 amplitudes than Match tones (Fig 3.C, t(18) = −4.10, p = 0.00068, Hedges’g = 0.42).

Together, these findings indicate that AI explainability and tone predictability independently modulated the P1–N1 complex. Whereas increasing levels of explanation were associated with a graded enhancement of sensory attenuation, mismatch tones consistently elicited larger P1–N1 amplitudes than match tones across condition.

N1–P2 Peak-to-Peak Amplitude

The analysis revealed significant effects of both AI Explainability and Tone Predictability on the N1–P2 complex. A main effect of AI Explainability was observed (F(2,36) = 4.55, p = 0.017, η²ₚ = 0.02), indicating that the type of explanation provided by the belief-based AI modulated the magnitude of the N1–P2 response.

Specifically, combined proximal + distal explanations significantly increased N1–P2 amplitudes relative to the no-explanation condition (post hoc: t(18) = 2.77, p = 0.0127, Hedges’ g = 0.35). In contrast, distal explanations alone did not differ significantly from the no-explanation condition (t(18) = 0.61, p = 0.55). The combined explanation condition also elicited significantly larger N1–P2 amplitudes than the distal-only condition (t(18) = −2.32, p = 0.0487, Hedges’ g = −0.265).

A robust main effect of Tone Predictability was also found (F(1,18) = 15.65, p = 0.00093, η²ₚ = 0.046), with mismatch tones (Fig. 3D) eliciting larger N1–P2 amplitudes than match tones (Fig. 3C; post hoc: t(18) = −3.96, p = 0.00093, Hedges’ g ≈ −0.45).

Finally, a trend-level interaction between AI Explainability and Tone Predictability was observed (F(2,36) = 2.94, p = 0.066), although this effect did not reach statistical significance after correction (p = 0.082, η²ₚ = 0.005).

Together, these findings indicate that AI explainability and tone predictability independently shaped the N1–P2 complex: combined explanations were associated with increased amplitudes, whereas mismatch tones consistently elicited larger responses than match tones across conditions.

##### MMN peak

To assess whether explanation richness modulated pre-attentive deviance detection, MMN peak amplitude was compared across the three explanation conditions (no explanation, distal explanation, combined proximal + distal explanation). Analyses revealed no significant effect of AI Explainability on MMN peak amplitude (F(2,36) = 0.097, p = 0.908, η²ₚ = 0.0029). This finding indicates that, consistent with Experiments 1 and 2, MMN responses remained stable across all explanation types, providing no evidence that explanation availability or richness influenced pre-attentive auditory deviance detection (see Figure 4.C).

## 4. Discussion

### 4.1. Impact of Automation on the Sense of Agency

Across behavioural and neural measures, automation produced a clear reduction in the SoA. This finding is consistent with a substantial body of literature showing that delegating control to automated systems weakens the perceived causal link between intention and outcome. The decrease in explicit FoC ratings aligns with previous work indicating that reduced motor involvement and increased system autonomy diminish metacognitive judgments of control (Berberian et al., 2012). Such effects are typically interpreted as reflecting a reduced contribution of reflective processes related to authorship and goal alignment when action selection is externally determined (Synofzik et al., 2013; Haggard & Chambon, 2012; Moore, 2016).

At the neural level, automation was associated with increased P1–N1 amplitudes, indicating reduced sensory attenuation when action selection was controlled by the AI rather than by the participant. Within predictive-processing frameworks, sensory attenuation is generally interpreted as reflecting the precision of predictions linking actions to their expected sensory consequences (Hughes et al., 2013; Haggard, 2017; Arnal & Giraud, 2012). When participants selected the vehicle’s destination themselves, these predictions were likely more precise, resulting in attenuated early auditory responses. Under automation, although participants still initiated the trial, they no longer determined the critical action outcome, namely the direction of the vehicle. This reduction in action–outcome contingency may have weakened predictive coupling, leading to larger P1–N1 responses.

In contrast, the N1–P2 complex showed reduced amplitudes under automation, with larger responses observed when participants retained control over action selection. This pattern is consistent with previous findings indicating that P2 modulation is sensitive not only to motor prediction but also to temporal parameters and task context. In the present paradigm, the action– outcome delay (300 ms) falls within a range in which P2 responses do not necessarily follow the classical suppression pattern typically observed for immediate action effects. Previous work has shown that P2 modulation is sensitive to action–outcome delays, with non-classical patterns, including enhanced responses for self-generated outcomes, observed when delays approach or exceed approximately 200 ms (Timm et al., 2016). More broadly, P2 activity has been associated with later stages of perceptual and evaluative processing (Crowley & Colrain, 2004). These results therefore suggest that automation influences not only early predictive processes but also intermediate stages of outcome processing. Within this framework, the larger N1–P2 amplitudes observed in the full-control condition suggest that later stages of outcome processing were more strongly engaged when participants retained control over action selection. By contrast, the smaller N1–P2 responses observed under automation indicate that these later stages of processing were modulated when control over action selection was externally delegated.

Taken together, these findings indicate that automation reduces the SoA at both behavioural and neural levels and affects multiple stages of action–outcome processing. Specifically, automation appears to weaken early predictive coupling, as reflected by reduced sensory attenuation within the P1–N1 complex, while also modulating later stages of outcome evaluation indexed by the N1–P2 complex.

### 4.2. Effects of Distal Explanations on Agency Restoration

Experiment 2 examined whether distal explanations could mitigate the loss of agency induced by automation. Behaviourally, distal explanations significantly increased the Feeling of Control (FoC), consistent with prior work showing that greater transparency and intelligibility of belief-based AI decisions enhance perceived control, trust, and alignment with automated systems (Vantrepotte et al., 2022; Houdoyer et al., under review). Providing information about the system’s higher-level goals may have allowed participants to reconstruct a more coherent link between their intentions and the system’s actions, thereby partially restoring explicit agency (Synofzik et al., 2013; Moore, 2016).

At the neural level, distal explanations were associated with reduced P1–N1 amplitudes relative to the no-explanation condition. This result is consistent with enhanced predictive processing, suggesting that distal information may have facilitated the formation of expectations about the system’s behaviour despite the absence of direct control over action selection (Arnal & Giraud, 2012; Hughes et al., 2013). This pattern is qualitatively consistent with the reduced P1–N1 amplitudes observed under voluntary control in Experiment 1. Nevertheless, because the two experiments were conducted in independent samples, this convergence should be interpreted cautiously and cannot be taken as direct evidence that distal explanations restore the neural signature of voluntary control.

Additionally, distal explanations were associated with an increase in the N1–P2 peak-to-peak complex. Interestingly, a similar pattern was observed in Experiment 1, where reduced P1–N1 and larger N1–P2 amplitudes were found in the condition associated with higher perceived control, namely the motor-control condition. This suggests that early (P1–N1) and intermediate (N1–P2) stages of outcome processing may be sensitive to factors related to perceived control under certain conditions. As noted above, later ERP components such as the P2 do not appear to reflect sensory prediction in a uniform manner. Their modulation depends on temporal and contextual factors, and may involve later evaluative or integrative stages of outcome processing rather than a simple attenuation mechanism (Timm et al., 2016; Pinheiro et al., 2019; Crowley & Colrain, 2004). Within this framework, the increase in the N1–P2 complex observed under distal explanations suggests that making the system’s higher-level intentions explicit modified later stages of outcome processing. More specifically, distal explanations may have made action outcomes easier to integrate within an interpretable action– outcome structure, thereby contributing to the observed increase in N1–P2 amplitudes.

Overall, these results suggest that distal explanations may partially restore the SoA by facilitating predictive engagement and influencing later stages of outcome processing, thereby contributing to a more coherent relationship between expected and observed system behaviour.

### 4.3. Effects of Combined Proximal and Distal Explanations

Experiment 3 showed that the strongest restoration of explicit agency occurred when proximal and distal explanations were combined. This pattern is consistent with theoretical accounts suggesting that users construct more accurate mental representations when explanations specify both why an action is taken and how it is implemented (Houdoyer et al., under review). By integrating goal-level and action-level information, combined explanations may have provided a more coherent representational framework, facilitating anticipation of system behaviour and strengthening perceived control.

At the neural level, this behavioural effect was accompanied by a graded modulation of ERP components. P1–N1 amplitudes decreased progressively from the no-explanation condition to distal explanations and further to combined explanations, consistent with increased predictive engagement as explanatory information became more complete. This effect was observed for both predicted and mispredicted outcomes, suggesting a general influence on anticipatory processing rather than a condition-specific effect. The N1–P2 peak-to-peak complex showed a complementary pattern, with larger amplitudes within the combined proximal and distal explanations configuration relative to either explanation condition taken in isolation.

Similar patterns were observed across Experiments 1 and 2, where attenuated P1–N1 and larger N1–P2 amplitudes were associated with conditions affording higher perceived control (the Motor condition in Experiment 1 and the belief-based AI with distal explanation condition in Experiment 2). In the present case, the larger N1–P2 amplitudes observed under combined explanations may indicate that providing both goal-level and action-level information strengthened later-stage processing of action outcomes beyond what was observed with distal explanations alone.

Overall, these findings indicate that explanatory richness may modulate multiple stages of action–outcome processing in a graded manner. By jointly specifying goal-level and action-level information, combined explanations appear to support more coherent hierarchical predictions, influencing both early anticipatory mechanisms and later stages of outcome integration. In this way, combined explanations may promote a more stable and interpretable SoA in Humain-AI interactions.

### 4.4. Effects of Tone Predictability Across Experiments

Across experiments, a significant main effect of Tone Predictability was observed, indicating that predicted and mispredicted tones were processed differently at the neural level. More specifically, predicted tones elicited a stronger attenuation of the P1–N1 complex compared to mispredicted tones. This pattern is consistent with predictive processing accounts, according to which predictable sensory events generate reduced neural responses due to lower prediction error (Arnal & Giraud, 2012; Hughes, Desantis, & Waszak, 2013). Within this framework, predicted tones likely benefited from more stable and precise expectations, whereas mispredicted tones, by violating these expectations, elicited larger neural responses.

Importantly, this effect was robust and consistent at the level of the P1–N1 complex across all experiments, whereas it was less stable for the N1–P2 complex, reaching significance only in Experiment 3. This dissociation suggests that early and later stages of auditory processing differ in their sensitivity to stimulus predictability. Specifically, the P1–N1 complex, which reflects early sensory processing, appears to rely on ERP components (i.e., P1 and N1) that provide the most robust index of predictive attenuation (Bendixen et al., 2012; Hughes et al., 2013). In contrast, the N1–P2 complex, associated with later stages of perceptual evaluation and contextual integration, appears more variable and more sensitive to the broader experimental context, coherently with previous findings of the P2 component (Crowley & Colrain, 2004; Timm et al., 2016; Pinheiro et al., 2019).

The pattern observed in Experiment 3 further supports this interpretation. In this experiment, a main effect of Tone Predictability on the N1–P2 complex was observed in the absence of any interaction with the degree of AI explanations (none, distal only, proximal + distal). This indicates that predictability effects generalized across all conditions, independently from the level of information provided by the belief-based AI.

Crucially, no interaction between Condition (Motor, belief-based AI without explanation, belief-based AI with distal explanation, and belief-based AI with distal + proximal explanations) and Tone Predictability was observed across experiments. This indicates that the difference between predicted and mispredicted tones was preserved across all action contexts, suggesting that the effect of tone predictability was largely independent of experimental manipulations related to control and explanation. Consistent with this view, Mismatch Negativity (MMN) waves were elicited across all experiments in response to mispredicted tones. However, MMN amplitudes did not differ significantly between conditions, indicating that the pre-attentive detection of auditory deviance remained stable across contexts (Näätänen et al., 2007).

Taken together, these findings suggest that tone predictability exerts a robust influence on early auditory prediction mechanisms, while its impact on intermediate stages of processing is more variable and dependent on the broader experimental context. The absence of interactions with Condition indicates that these effects are largely independent of the manipulations of control and explanation, pointing to a relatively stable influence of sensory predictability across agency contexts.

### 4.5. Convergence Between Behavioural and Neural Markers of Agency

Across the three experiments, behavioural and electrophysiological findings revealed a coherent and interpretable pattern. Automation consistently reduced the SoA at both subjective and neural levels, whereas explanatory information was associated with parallel changes in these measures. The strongest effects were observed when proximal and distal explanations were combined, suggesting that explanatory richness may support agency through coordinated influences on reflective evaluations and predictive processing.

As detailed in the Materials and Methods, explicit agency ratings were collected on a subset of trials and were not intended to support trial-level correlations with electrophysiological measures. Rather, they served to confirm that the experimental manipulations reliably affected subjective experience. In contrast, electrophysiological analyses were conducted across the full set of trials and revealed condition-dependent modulations of auditory ERP components, including P1–N1 and N1–P2 peak-to-peak measures.

Although behavioural and neural measures cannot be directly mapped onto one another, their consistent variation across conditions supports an integrated interpretation in which explanatory information influences both subjective agency and the processing of action outcomes.

This pattern is consistent with hierarchical accounts of agency, according to which the sense of control emerges from the interaction between predictive mechanisms and higher-level inferential processes (Chambon et al., 2011, 2017; Synofzik et al., 2013; Haggard, 2017).

Across all experiments, MMN responses remained stable, indicating that the automatic detection of auditory deviance (Näätänen et al., 2007) was not measurably affected by automation or explanation structure and was always observable. By contrast, the modulation of early and intermediate ERP complexes suggests that the observed effects primarily involve processes related to prediction and outcome processing, rather than more general mechanisms of auditory deviance detection.

### 4.6. Implications for Explainable AI and Human-AI Interaction

The present findings carry important implications for the design of explainable AI (XAI) systems. They indicate that explainability is not only associated with explicit evaluations such as trust, understanding, or perceived usability, but also with modulations of neurocognitive processes involved in the anticipation and processing of system behaviour. In this respect, explanatory information appears to operate across multiple levels of processing, influencing both reflective evaluations and predictive engagement.

The particularly strong effects observed for combined explanations highlight the potential importance of multi-level explanatory structures. Explanations that jointly specify why an action is performed and how it unfolds may provide a more coherent representational framework, facilitating the formation of expectations about system behaviour. This interpretation is consistent with human-centred approaches to XAI, which emphasise context-sensitive explanation strategies (Miller, 2019).

More broadly, these findings suggest that effective explainability should not be conceived solely as information provision, but also as a means of supporting alignment between user expectations and system behaviour. This perspective is consistent with frameworks that conceptualise explainability as supporting user understanding and appropriate interaction with automated systems (Miller, 2019). By influencing both early predictive processes and later stages of outcome processing, explanatory information may help maintain a coherent SoA even in contexts where control is partially delegated.

Incorporating insights from cognitive neuroscience may therefore offer a valuable perspective for the design of XAI systems. Explanation strategies that support the formation of coherent internal representations of system behaviour may contribute to sustaining user engagement and agency in increasingly automated environments.

## 5. Conclusion and Perspectives

Across three experiments, automation was associated with a reduction in both explicit and neural correlates of the SoA, whereas explanations, particularly those combining proximal (path-level) and distal (goal-level) information, were associated with graded and component-specific changes in these markers. Modulations of P1–N1 and N1–P2 complexes suggest that explanatory information influences predictive engagement and intermediate stages of outcome processing. The stability of the MMN across all manipulations indicates that the automatic detection of auditory deviance was not measurably affected by automation or explanation structure.

These results identify a set of neurocognitive correlates of agency-related processing that can be measured without interrupting behaviour, which may be relevant for the study of agency in applied contexts such as autonomous driving. They also suggest that adaptive AI systems could, in principle, benefit from adjusting explanation formats or control-sharing strategies in response to users’ cognitive states, although such applications remain to be empirically validated.

Beyond discrete ERP measures, the present findings motivate extending the study of agency to more continuous and dynamic forms of interaction. In naturalistic settings such as driving, neural responses overlap and evolve over time, which limits the applicability of single-event ERP analyses.

Oscillatory markers such as alpha–mu event-related desynchronization (ERD), which have been linked to motor preparation and attentional processes (Pfurtscheller & Lopes da Silva, 1999; Engel & Fries, 2010), have also been associated with variations in perceived control in continuous tasks (Wen et al., 2017; Kang et al., 2015). Future work combining such oscillatory features with data-driven approaches, including deep learning models applied to continuous EEG signals, may provide new opportunities to track evolving agency-related states in ecologically valid human–AI interaction contexts.

## References

Arnal, L. H., & Giraud, A.-L. (2012). Cortical oscillations and sensory predictions. Trends in Cognitive Sciences, 16(7), 390–398.

Baess, P., & others. (2011). Selective suppression of self-initiated sounds in an auditory stream: An ERP study. Psychophysiology, 48(9), 1276–1283.

Bauer, K., Hinz, O., van der Aalst, W., & Weinhardt, C. (2021). Expl(AI)n it to me – Explainable AI and information systems research. Business & Information Systems Engineering, 63(2), 79–82.

Bednark, J. G., Poonian, S. K., Palghat, K., McFadyen, J., & Cunnington, R. (2015). Identity-specific predictions and implicit measures of agency. *Psychology of Consciousness: Theory*, Research, and Practice, 2, 253–268.

Bendixen, A., SanMiguel, I., & Schröger, E. (2012). Early electrophysiological indicators for predictive processing in audition: A review. International Journal of Psychophysiology, 83, 120–131.

Berberian, B., Sarrazin, J. C., Le Blaye, P., & Haggard, P. (2012). Automation technology and sense of control: A window on human agency. PLoS ONE, 7(3), e34075.

Bigenwald, A., & Chambon, V. (2019). Criminal responsibility and neuroscience: No revolution yet. Frontiers in Psychology, 10, Article 1406.

Bolt, N. K., & Loehr, J. D. (2021). Sensory attenuation of the auditory P2 differentiates self-from partner-produced sounds during joint action. Journal of Cognitive Neuroscience, 33, 2297–2310.

Chambon, V., Domenech, P., Jacquet, P. O., Barbalat, G., Bouton, S., Pacherie, E., Koechlin, E., & Farrer, C. (2017). Neural coding of prior expectations in hierarchical intention inference. Scientific Reports, 7, Article 1278.

Chambon, V., Domenech, P., Pacherie, E., Koechlin, E., Baraduc, P., & Farrer, C. (2011). What are they up to? The role of sensory evidence and prior knowledge in action understanding. PLoS ONE, 6(2), e17133.

Christoffersen, K., & Woods, D. D. (2002). How to make automated systems team players. In Advances in Human Performance and Cognitive Engineering Research. Emerald Group Publishing.

Crowley, K. E., & Colrain, I. M. (2004). A review of the evidence for P2 being an independent component process: Age, sleep and modality. Clinical Neurophysiology, 115(4), 732–744.

Duzcu, H., Özkurt, T. E., Mapelli, I., & Hohenberger, A. (2019). N1–P2: Neural markers of temporal expectation and response discrimination in interval timing. Acta Neurobiologiae Experimentalis, 79(2), 193–204.

Egan, S., Weber, C., Ghio, M., & Bellebaum, C. (2026). I, You, Robot: Attenuation for auditory outcomes of actions performed by different agents shows distinct patterns for N1 and P2 amplitudes. Biological Psychology, Article 109241.

Endsley, M. R., & Kiris, E. O. (1995). The out-of-the-loop performance problem and level of control in automation. Human Factors, 37(2), 381–394.

Getzmann, S., Wascher, E., & Schneider, D. (2018). The role of inhibition for working memory processes: ERP evidence from a short-term storage task. Psychophysiology, 55(5), e13026.

Haggard, P. (2017). Sense of agency in the human brain. Nature Reviews Neuroscience, 18, 197–208.

Han, N., Jack, B. N., Hughes, G., Elijah, R. B., & Whitford, T. J. (2021). Sensory attenuation in the absence of movement: Differentiating motor action from sense of agency. Cortex, 141, 436–448.

Houdoyer, E., Berberian, B., Le Bars, S., & Chambon, V. (under review). Aligning minds and machines: Hierarchical explanations enhance sense of agency in AI-assisted decision-making. Research Square.

Hughes, G., Desantis, A., & Waszak, F. (2013). Attenuation of auditory N1 responses to self-generated sounds: Evidence for internal forward models. Psychological Bulletin, 139(1), 133–151.

Hughes, G., Desantis, A., & Waszak, F. (2013). Attenuation of auditory N1 results from identity-specific action-effect prediction. European Journal of Neuroscience, 37(7), 1152– 1158.

Kühn, S., Nenchev, I., Haggard, P., Brass, M., Gallinat, J., & Voss, M. (2011). Whodunnit? Electrophysiological correlates of agency judgements. PLoS ONE, 6, e28657.

Kuznetsova, A., Brockhoff, P. B., & Christensen, R. H. B. (2017). lmerTest package: Tests in linear mixed effects models. Journal of Statistical Software, 82, 1–26.

Le Bars, S., Bourgeois-Gironde, S., Wyart, V., Sari, I., Pacherie, E., & Chambon, V. (2022). Motor coordination and strategic cooperation in joint action. Psychological Science.

Le Bars, S., Darriba, Á., & Waszak, F. (2019). Event-related brain potentials to self-triggered tones: Impact of action type and impulsivity traits. Neuropsychologia, 125, 14– 22.

Le Bars, S., Devaux, A., Nevidal, T., Chambon, V., & Pacherie, E. (2020). Agents’ pivotality and reward fairness modulate sense of agency in cooperative joint action. Cognition, 195, Article 104117.

Loehr, J. D. (2013). Sensory attenuation for jointly produced action effects. Frontiers in Psychology, 4, Article 172.

Luke, S. G. (2017). Evaluating significance in linear mixed-effects models in R. Behavior Research Methods, 49(4), 1494–1502.

Moore, J. W., & Obhi, S. S. (2012). Intentional binding and the sense of agency: A review. Consciousness and Cognition, 21(1), 546–561.

Näätänen, R., Kujala, T., & Winkler, I. (2011). Auditory processing that leads to conscious perception: A unique window to central auditory processing opened by the mismatch negativity and related responses. Psychophysiology, 48(1), 4–22.

Pacherie, E. (2008). The phenomenology of action: A conceptual framework. Cognition, 107(1), 179–217.

Pagliari, M., Chambon, V., & Berberian, B. (2022). What is new with artificial intelligence? Human–agent interactions through the lens of social agency. PsyArXiv.

Pinheiro, A. P., Schwartze, M., Gutierrez, F., & Kotz, S. A. (2019). When temporal prediction errs: ERP responses to delayed action-feedback onset. Neuropsychologia, 134, Article 107200.

Putnam, V., & Conati, C. (2019). Exploring the need for explainable artificial intelligence (XAI) in intelligent tutoring systems. In IUI Workshops.

Ross, B., Barat, M., & Fujioka, T. (2017). Sound-making actions lead to immediate plastic changes of neuromagnetic evoked responses and induced β-band oscillations during perception. Journal of Neuroscience, 37(24), 5948–5959.

Sarter, N. B., Woods, D. D., & Billings, C. E. (1997). Automation surprises. In Handbook of Human Factors and Ergonomics (pp. 1926–1943).

Sugimoto, F., Kimura, M., & Takeda, Y. (2021). Attenuation of auditory N2 for self-modulated tones during continuous actions. Biological Psychology, 166, Article 108201.

Synofzik, M., Vosgerau, G., & Voss, M. (2013). The experience of agency: An interplay between prediction and postdiction. Frontiers in Psychology, 4, Article 127.

Timm, J., SanMiguel, I., Keil, J., Schröger, E., & Schönwiesner, M. (2014). Motor intention determines sensory attenuation of brain responses to self-initiated sounds. Journal of Cognitive Neuroscience, 26, 1481–1489.

Timm, J., Schönwiesner, M., Schröger, E., & SanMiguel, I. (2016). Sensory suppression of brain responses to self-generated sounds is observed with and without the perception of agency. Cortex, 80, 5–20.

van der Wel, R. P., Sebanz, N., & Knoblich, G. (2012). The sense of agency during skill learning in individuals and dyads. Consciousness and Cognition, 21(3), 1267–1279.

Vantrepotte, Q., Berberian, B., Pagliari, M., & Chambon, V. (2022). Leveraging human agency to improve confidence and acceptability in human–machine interactions. Cognition, 222, Article 105020.

Wen, W., Yamashita, A., & Asama, H. (2017). Measurement of the perception of control during continuous movement using electroencephalography. Frontiers in Human Neuroscience, 11, Article 392.

Widmann, A., & Schröger, E. (2022). Intention-based predictive information modulates auditory deviance processing. Frontiers in Neuroscience, 16, Article 995119.

